# Efficacy of Minnelide in a Next-Generation Dual-Recombinase Regulated Genetically Engineered Mouse Model of CIC::DUX4 Sarcoma

**DOI:** 10.1101/2025.11.06.687065

**Authors:** MaKenna R. Browne, Axel V. Silver, Risha Banerjee, Benigno Aquino, Kristianne M. Oristian, Jonathon E. Himes, Peter G. Hendrickson, David G. Kirsch

## Abstract

CIC::DUX4 sarcoma (CDS) is a lethal cancer driven by a fusion between tumor suppressor Capicua (CIC) and pioneer transcription factor double homeobox 4 (DUX4). To develop an immunocompetent pre-clinical model of CDS, we previously generated three genetically engineered mouse models (GEMMs) of CDS with *CIC::DUX4* regulated by *loxP*-STOP-*loxP* cassettes. However, all three models developed spontaneous tumors without Cre recombinase. Here, we established an innovative GEMM of CDS (dFLEx CDS) that employs a dual recombinase (Cre + FLPE) FLEx-switch design to activate *CIC::DUX4* expression and initiate sarcomagenesis in a spatially and temporally-controlled manner. Because CIC::DUX4 drives sarcoma development by activating a distinct oncogenic transcriptional program, we performed a drug screen on human-derived CDS cell lines using a library of compounds that modulate transcriptional regulation. This screen identified Minnelide, an inhibitor of RNA polymerase II-mediated transcription, as a selective inhibitor of CDS. Mechanistically, Minnelide acted through xeroderma pigmentosum type B to alter phosphorylation of RPB1, the largest subunit of RNA polymerase II. Subsequently, RPB1 underwent degradation leading to apoptosis of CDS cells. Minnelide demonstrated in vivo efficacy in autochthonous dFLEx CDS GEMMs and in human CDS xenografts. As Minnelide has already been demonstrated to be safe in clinical trials with activity for adult cancers, these findings nominate Minnelide as a novel therapeutic option to test in CDS patients.

## Introduction

CIC::DUX4 sarcoma (CDS) is a soft tissue sarcoma that typically affects adolescents and young adults and is driven by t(4;19) or t(10;19) chromosomal translocations, which is the most common genetic rearrangement found in EWSR1 fusion-negative small round blue cell sarcomas (1–4). Until recently, CDS was referred to as “Ewing-like sarcoma” due to similar histological and immunohistochemical features of Ewing sarcoma. However, unlike Ewing sarcoma, CDS 1) rarely arises in the bone, 2) lacks the Ewing-defining ETS gene fusion, and 3) is associated with a more aggressive clinical course. Although the World Health Organization recently classified CIC-rearranged sarcomas as a distinct entity (5), patients are still routinely treated with Ewing sarcoma-based chemotherapy despite mounting data demonstrating limited efficacy, high rates of recurrence, and poor outcomes (2,3,6). Therefore, novel treatments that are effective in CDS are urgently needed.

Mechanistically, CDS is driven by a unique fusion protein composed of the majority of the transcriptional repressor Capicua (CIC), including its DNA binding domain, and the C-terminal transactivation domain of the pioneer transcription factor double homeobox 4 (DUX4)(7). Endogenous CIC represses the expression of genes downstream of receptor tyrosine kinase (RTK) activation (8–11), however, upon fusion to the C-terminal transactivation domain of DUX4, CIC::DUX4 functions as a potent transcriptional activator at normally repressed CIC binding sites (7,12–14). Given CDS is fundamentally driven by activation of an aberrant transcriptional program, modulating transcription could serve as a promising therapeutic approach.

A significant barrier to identifying better therapeutic options for CDS, is the lack of robust pre-clinical models for identifying and testing novel therapies. While many studies have successfully utilized patient-derived and ectopically-derived xenograft models (12,13,15–19) to interrogate key oncogenic mechanisms of CDS, these models are limited by the absence of an intact immune system and the natural tumor microenvironment which is critical for tumor growth and response to therapy. Previous work reported on the development of a transgenic CIC::DUX4 sarcoma Zebrafish model (20), however, generation of an immunocompetent genetically engineered mouse model (GEMM) has been challenging. Our first attempts to generate GEMMs used *loxP*-STOP-*loxP* cassettes and relied on the expression of Cre recombinase to activate the expression of CIC::DUX4 either at the *Rosa26* locus or at the endogenous *Cic* locus (21). Notably, in the absence of Cre, all three models developed CDS tumors spontaneously and progressed rapidly before the mice were able to breed preventing colony maintenance (21). Collectively, these experiments demonstrated that CDS could be modeled in immunocompetent murine systems, however novel approaches were necessary to spatially and temporally-control CIC::DUX4 expression.

In this study, we developed an innovative GEMM of CDS (dFLEx CDS) that uses two independent recombinases (Cre + FLPE) to activate *CIC::DUX4* expression in a controlled manner. Furthermore, we conducted a drug screen on human-derived CDS cell lines using a library of compounds that modulate transcription including drugs that selectively target a diverse array of epigenetic writers and erasers. By leveraging our *in vitro* drug study, human CDS xenografts, and novel dFLEx CDS GEMM, we demonstrate that Minnelide, an inhibitor of RNA polII transcription, is a promising therapeutic approach for CDS.

## Results

### Dual recombinase regulation of CIC::DUX4 expression results in a spatially and temporally-restricted CDS mouse model

A significant barrier to pre-clinical investigation of novel therapeutic approaches for CDS is the lack of an immunocompetent, autochthonous mouse model. Our lab previously attempted to develop a GEMM of CDS by targeting mouse embryonic stem (ES) cells using three different approaches that each regulated CIC::DUX4 expression with *loxP*-STOP-*loxP* cassettes (21). Remarkably, mice derived from the ES cells in all three models spontaneously developed aggressive and multi-focal sarcomas in the absence of Cre recombinase leading to rapid and early animal demise (21). Therefore, in these models we were unable to control the timing or location of tumor formation and it was not possible to breed the transgenic allele. To address these limitations, we developed a fourth CDS mouse model in which CIC::DUX4 expression requires both Cre and FLP recombinases to conditionally invert two exons into the correct orientation and reading frame (**Figure 1A**). In the absence of Cre and FLP recombinases, a neomycin STOP cassette precedes an inverted exon 1 (CIC), SV40pA-KT3 stuffer sequence (pA), and inverted exon 2 (DUX4). Cre-mediated recombination inverts exon 1 into the same orientation as the CAG promoter, and excises the neomycin STOP cassette. Following Cre recombination, the SV40pA-KT3 stuffer sequence remains intact, and the inverted exon 2 is out of frame. FLP-mediated recombination excises the SV40pA-KT3 stuffer sequence and inverts exon 2 into the same orientation and in frame with the remainder of the *CIC::DUX4* transcript. Both Cre and FLP-mediated recombination are therefore required for *CIC::DUX4* transgene expression (**Figure 1A)**. Indeed, in the absence of Cre and Flp, the dFLEx CDS system prevented spontaneous tumor formation. Following electroporation of pCAG-Cre and pCAG-

**Figure 1.**
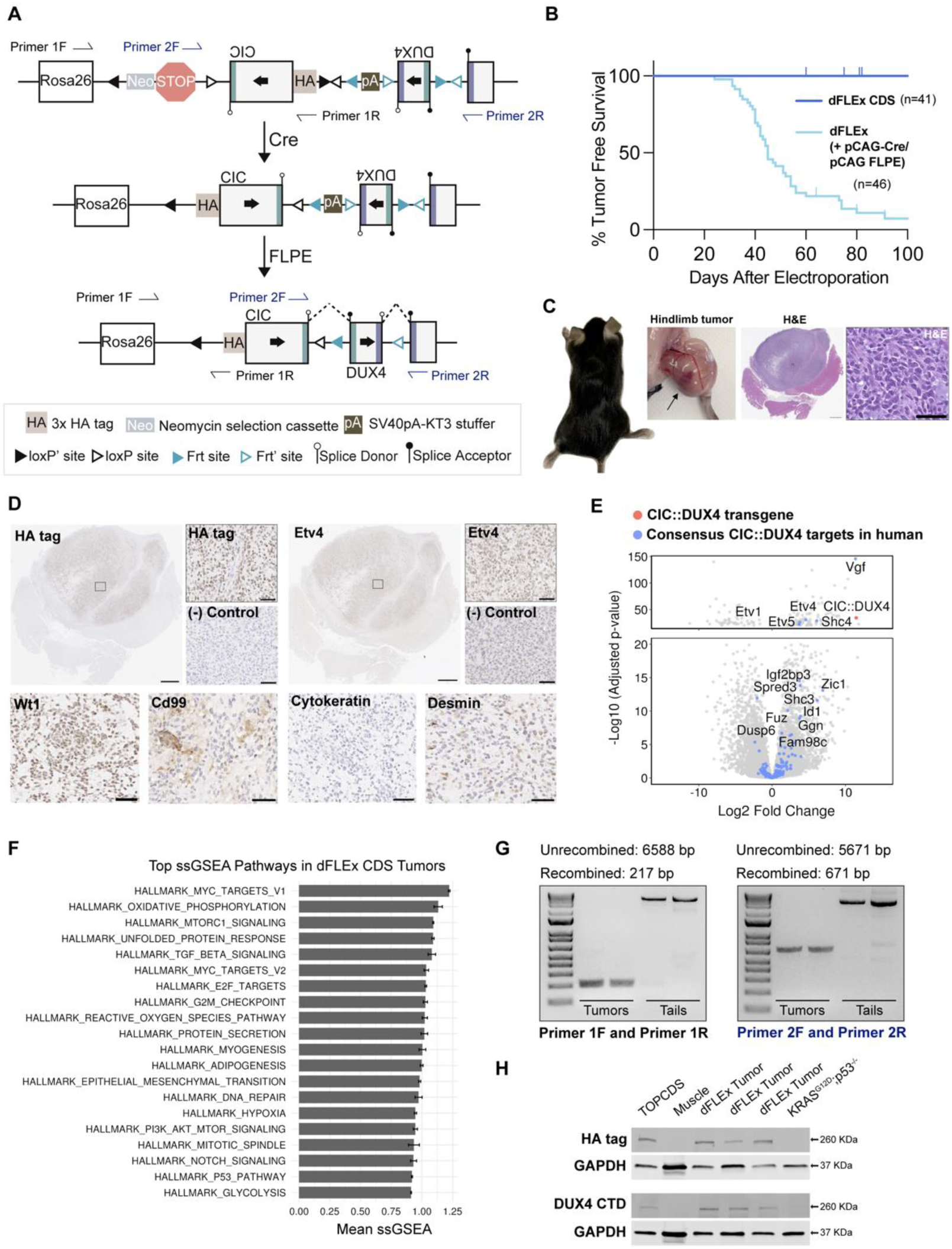
Dual-Flex (dFLEx) CDS mice develop tumors only after expression of Cre and FLPE. **A**. Schematic of the dFLEx CDS allele before and after recombination with Cre and FLPE. PCR primer sequences used to assess recombination are indicated with Primer 1F/R and Primer 2F/R arrows. **B.** Kaplan-Meier survival curve of dFLEx CDS mice showing no spontaneous tumor development without Cre and FLPE and ∼90% tumor penetrance after electroporation of the hindlimb muscle with plasmids expressing Cre + FLPE. **C.** Gross images of a hindlimb tumor and H&E stained sections of a soft tissue CDS tumor. Scale bars 1500 μm (left) and 50μm (right). **D.** Immunohistochemistry on tumors formed in dFLEx CDS mice after electroporation of Cre + FLPE plasmids for HA tag and the indicated proteins. Sarcomas from KRAS^G12D^; p53^fl/fl^ (KP) mice used as negative (-) control. Scale bars 1500 μm (entire tumor sections) and 50μm (insets denoted by boxes). E. Volcano plot demonstrating strong induction of CIC::DUX4 consensus transcriptional target genes in dFLEx CDS tumors relative to KP control. **F.** GSEA pathway analysis on dFLEx CDS tumors **G.** DNA gel of amplification products from PCR of genomic DNA from tumors and tails (n=2 each) using primers indicated in the schematic in **A** that span the loxP and FRT sites. In the tumors a ∼217bp and ∼671bp product is amplified consistent with recombination. **H.** Western blot on lysates from dFLEx CDS tumors probed for the HA-tag and DUX4 CTD confirming the expression of a ∼ 260kD protein in dFLEx CDS tumors (n=3). TOPCDS mouse CDS cell line (Hendrickson et al. Oncogene 2024) was used as a positive control and dFLEx CDS contralateral muscle and a KP cell line were used as negative controls.

FLPE plasmids into the hindlimbs of heterozygous and homozygous dFLEx mice, spatially restricted tumors formed within 3 months at ∼90% penetrance (**Figure 1B and C**). Hematoxylin and eosin staining of sections from these tumors exhibited the expected small round blue cell morphology (**Figure 1C)**, with strong HA tag, Etv4, Wt1, weak Desmin expression, as well as focal/patchy Cd99 expression, but not Cytokeratin consistent with human CIC::DUX4 fusion-positive tumors (**Figure 1D**). Furthermore, RNA sequencing comparing the transcriptional profiles of the dFLEx CDS tumors compared to sarcomas in KRAS^loxP-STOP-loxP-G12D^ p53^fl/fl^; (KP) mice, demonstrates dFLEx CDS tumors express known CIC::DUX4 target genes (14,21) such as *Etv1/4/5*, *Dusp6*, *Shc3/4*, and *Spred3* (**Figure 1E**). Additionally, GSEA analysis indicates enrichment of several notable driver oncogenic signaling pathways including Myc targets, Mtor, Notch, PI3k-Akt, and Tgfβ signaling (**Figure 1F).** PCR amplification across the *loxP* and FRT sites confirmed successful recombination by Cre and FLPE respectively (**Figure 1G**), The CIC::DUX4 fusion protein was detected in dFLEx tumors by western blot with antibodies for HA tag and DUX4 (**Figure 1H**). Collectively, these results demonstrate that the use of two independent recombinase systems prevents spontaneous tumor formation in the absence of a recombinase, while electroporation of separate pCAG-Cre and pCAG-FLPE plasmids enables spatially and temporally-restricted sarcomas that mimic human CDS.

### Minnelide kills CIC::DUX4 sarcoma cells through induction of apoptosis

Following the development of the dFLEx CDS model, we aimed to identify and evaluate novel therapeutic interventions for CDS using this platform. Considering CIC::DUX4 interacts with acetyltransferase p300 to activate a unique oncogenic transcriptional program (14,16), we sought to test the efficacy of compounds that modulate transcription or selectively target epigenetic writers and erasers. To this end we performed cell viability screens in 3 human CDS cell lines (Kitra-SRS, CDS2, X1C1) at 4 concentrations (1nm, 10nm, 100nm and 1000nM) using a library of 160 small molecule inhibitors (**Figure 2A, Supplemental Figure 1**). This drug screen not only identified Dinaciclib, a transcriptional modulator previously shown to have efficacy in CDS models (12), but also revealed a novel sensitivity to Triptolide even at the low nanomolar range tested (**Figure 2A).** Although Triptolide has demonstrated robust anti-tumor effects in a variety of pre-clinical cancer models (22–32), the clinical utility of Triptolide is limited by its poor solubility in water. Therefore, a water-soluble prodrug of Triptolide called Minnelide was developed (33). Minnelide rapidly releases Triptolide when exposed to phosphatases present in both tissues and in the blood (33). Importantly, Minnelide has been successfully tested in Phase I and II clinical trials for advanced gastrointestinal (GI) carcinoma and pancreatic cancer (34,35) and is currently being studied in ongoing clinical trials of gastric cancer (NCT05566834) and small cell lung cancer (NCT05166616). To validate our screen results and to test the efficacy of Minnelide in CDS cells relative to other small round cell sarcoma cell lines, we performed CellTiter-Glo assays. After 48h of treatment, CDS cells (Kitra-SRS, ECD1, 690108) exhibited reduced viability compared to Ewing sarcoma (A-673), fusion-positive rhabdomyosarcoma (Rh-4) and wild-type mouse embryonic fibroblasts (WT MEFs) (**Figure 2B**). Previous studies have shown Minnelide/Triptolide induces apoptosis in some cancer cell types, and autophagy in others (25–27,30,31,36–39). To explore the mechanism of cell death in CDS cells, we treated Kitra-SRS and ECD1 human CDS cells with 25nM Minnelide for 48 hours, and then performed Annexin V/ propidium iodide flow cytometry. In both cell lines tested, Minnelide treatment resulted in an increase of Annexin V^+^/ propidium iodide^-^ cells indicating early apoptosis (**Figure 2C, Supplemental Figure 2A**). Additionally, after 48h, there was an increase in cleaved-caspase 3 expression (**Figure 2D**) and a decrease in EdU incorporation (**Supplemental Figure 2B**) further suggesting Minnelide induces apoptosis in CDS cells. To begin to explore how Minnelide might induce apoptosis, we analyzed changes in the expression of genes that regulate the intrinsic and extrinsic pathway of apoptosis over 72 hours. Minnelide treatment increased the expression of the proapoptotic gene BCL2L11, which encodes for the BH3-only protein BIM and decreased the expression of the anti-apoptotic gene BCL2 (**Figure 2E**), suggesting that Minnelide may activate the intrinsic pathway of apoptosis.

**Figure 2.**
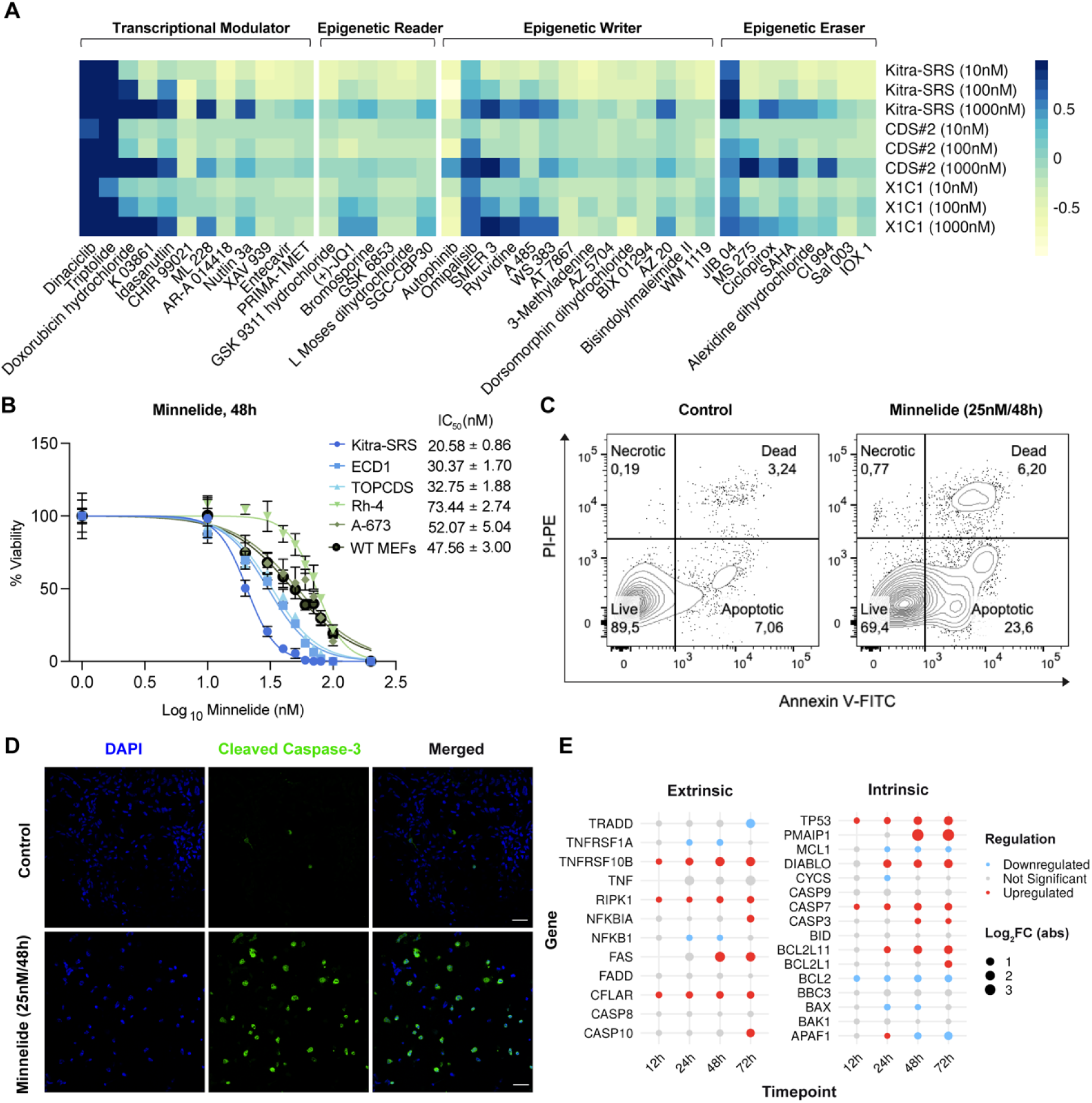
Minnelide induces apoptosis in CDS cells. **A**. Viability drug screen on human CDS cells using the Tocriscreen Epigenetics 3.0 compound library. **B.** Cell-titer Glo on CDS (Kitra-SRS, ECD1, TOPCDS mouse CDS cell line (Hendrickson et al. Oncogene 2024)) and Non-CDS (Rh-4, A-673, WT MEFs) cells treated with 0-200nM Minnelide for 48h. **C.** Annexin V/PI flow cytometry on Kitra-SRS CDS cells treated with Minnelide for 48h. **D.** Cleaved-Caspase 3 immunofluorescence on human CDS cells treated with Minnelide for 48h. **E.** Expression of apoptotic regulatory genes in Kitra-SRS cells 12, 24, 48 and 72h after treatment with 25nM Minnelide based on RNA-sequencing.

### Minnelide directly targets XPB leading to RPB1 degradation and inhibition of transcription

Previous studies have shown that Minnelide/Triptolide directly binds to xeroderma pigmentosum type B (XPB), a subunit of Transcription factor II H (TFIIH), which is a general transcription factor of RNA polymerase II (RNAP II) (28,40–42). In this working model, upon binding, Minnelide inhibits XPB’s ATPase activity leading to stalling of RNAP II at gene promoters and inhibition of sustained transcription. Prolonged stalling of RNAP II can result in altered phosphorylation patterns on RPB1, the largest subunit of RNAP II, ubiquitination of RPB1, followed by proteasomal mediated degradation of RPB1 (40–42) (**Figure 3A**). Consistent with this model, after treating human CDS cells (Kitra-SRS) with 10 μM Minnelide for 10-180 minutes, we observed an initial increase in RPB1 phosphorylation at Ser5 (marking transcriptional initiation) while RPB1 phosphorylation at Ser2 (marking transcriptional elongation) was maintained. By 120 minutes, total RPB1 levels were markedly reduced (**Figure 3B**). To test if the RPB1 reduction was due to proteasomal mediated degradation, CDS cells were co-treated with 10 μM Minnelide and increasing concentrations of proteosome inhibitor Epoxomicin. Co-treatment with Epoxomicin rescued the Minnelide-mediated decrease in RPB1 expression (**Figure 3C**), suggesting Minnelide treatment leads to the proteasomal-mediated degradation of RPB1 in CDS cells. Next, to test the model that Minnelide regulates RPB1 degradation via XPB, a Minnelide resistant, XPB mutant (42) (XPB C342T) was expressed in human CDS cells (**Figure 3D**). Human CDS cells expressing XPB C342T not only retained RPB1 expression after 48h Minnelide treatment, (**Figure 3E)** but also demonstrated a higher tolerance to Minnelide across increasing concentrations (**Figure 3F, Supplemental Figure 2A**). Taken together, these results suggest that Minnelide targets XPB leading to RNAP II stalling and subsequent RPB1 degradation, thus inhibiting CDS cell viability.

**Figure 3.**
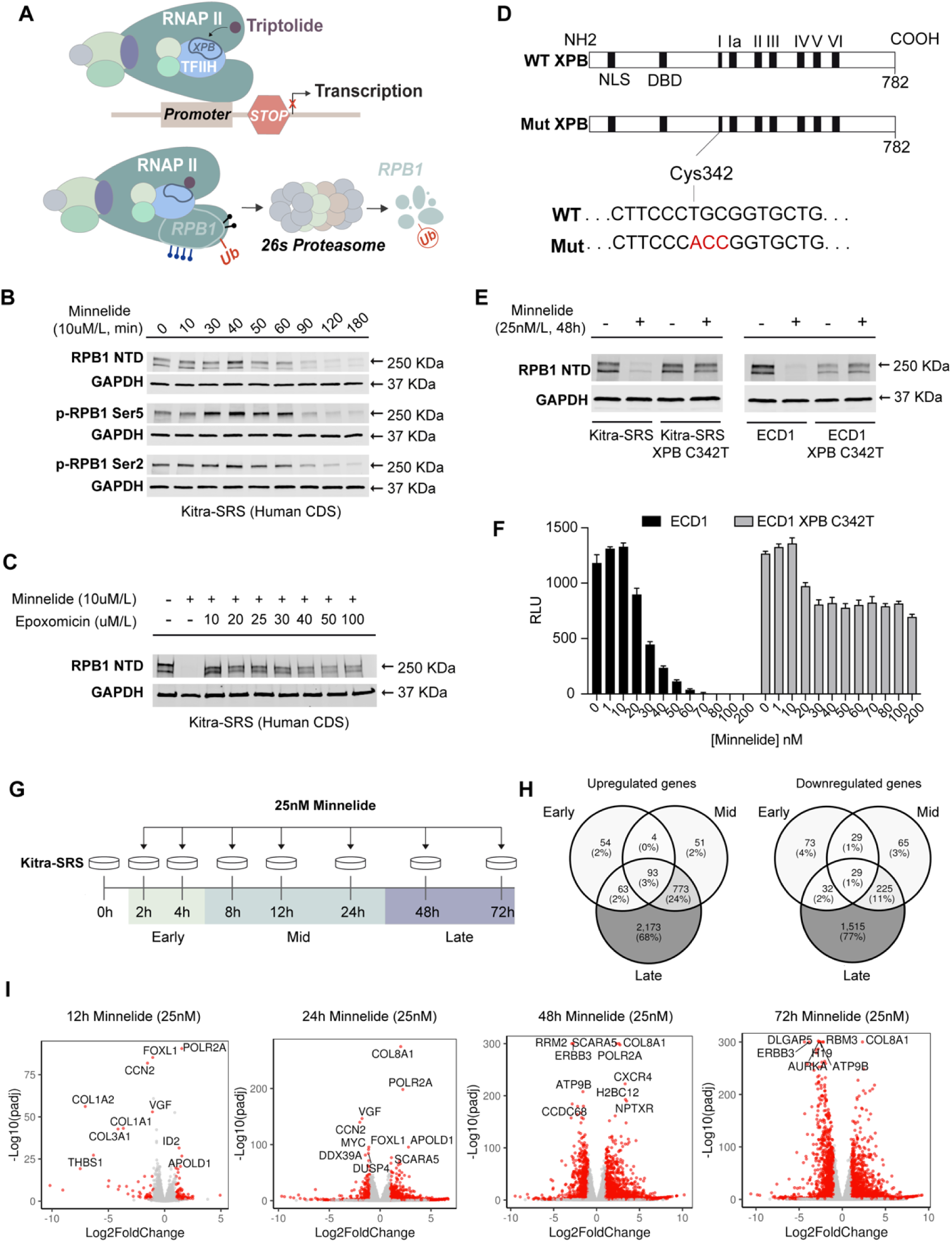
Minnelide targets XPB leading to RPB1 degradation and transcriptional inhibition. **A.** Schematic for Minnelide inhibition of RNAP II. Minnelide directly binds xeroderma pigmentosum type B (XPB). Inhibition of XPB’s ATPase activity leads to stalling of RNAP II at gene promoters, and inhibition of transcription. Prolonged stalling of RNAP II results in altered phosphorylation patterns on RPB1, ubiquitination, then proteasomal mediated degradation of RPB1. **B.** Western blot of RPB1 and phosphorylated RPB1 over a time course of Minnelide treatment in human Kitra-SRS CDS cells. **C.** Western blot demonstrating proteosome inhibitor Epoxomicin rescues Minnelide mediated RPB1 degradation in Kitra-SRS CDS cells. **D.** Strategy for generating a Minnelide-resistant XPB mutant. **E.** Western blot demonstrating expression of XPB C342T in Kitra-SRS and ECD1 human CDS cells rescues Minnelide mediated RPB1 degradation. **F.** Cell-titer-glo assay demonstrates human ECD1 CDS cells expressing XPB C342T have increased resistance to Minnelide. **G.** Schematic for Minnelide time course in Kitra-SRS human CDS cells. **H.** Venn diagrams representing transcriptional changes during early, mid and late Minnelide treatment times. **I.** Volcano plots demonstrating transcriptional changes following Minnelide treatment as a function of time.

To investigate the mechanism by which RNAP II stalling and transcriptional inhibition impacts CDS viability, we treated human CDS cells (Kitra-SRS) with 25nM Minnelide for 2, 4, 8, 12, 24, 48 and 72h, and then conducted bulk RNA sequencing. Previous studies in G3 medulloblastoma (G3 MB) demonstrated that Minnelide-induced cell death occurs, in part, through an early reduction of the G3 MB dependency gene, *MYC* (28). Considering these findings and the known function of CIC::DUX4 as a neomorphic transcriptional activator, we hypothesized that the effect of Minnelide on viability would be conferred through the altered expression of key oncogenic CIC::DUX4 target genes, such as *ETV1/4/5*, at early to mid-treatment timepoints **(Figure 3G**). Surprisingly, only a subset of CIC::DUX4 target genes emerged as top downregulated genes following 2-24h of Minnelide treatment. Target genes such as *DUSP4*, *VGF* and *MYC* were significantly downregulated at 12 and 24h (**Figure 3I**), however, the majority of CDS target genes including *ETV1/4/5,* remained relatively unchanged until 72h (**Supplemental Figure 4)**. These results suggest Minnelide-induced cell death could be occurring through a subset of downregulated CIC::DUX4 target genes alone or in combination with other essential genes (e.g. MYC targets) which are preferentially impacted by Minnelide.

### Minnelide treatment reduces tumor growth in a subset of dFLEx CDS mice

To test the efficacy of Minnelide against CDS in an immunocompetent, autochthonous setting, pCAG-Cre and pCAG-FLPE plasmids were electroporated into the hindlimb muscle of dFLEx CDS mice. Once tumors were palpable (∼40 days post electroporation), mice were treated daily for 21 days with either 0.42mg/kg Minnelide, 0.27mg/kg Minnelide or vehicle control (saline) (**Figure 4A**). Dose selection in this mouse study was informed by previous pre-clinical studies using Minnelide (28,33,43), and the maximum tolerated dose reported in the Phase I Minnelide clinical trial for gastrointestinal cancers (34). A dose of 0.27mg/kg of Minnelide approximates 0.80 mg/m^2^ in human patients (44) which has been successfully used in the Phase I Minnelide clinical trial for gastrointestinal cancers (34). Treatment with both 0.42mg/kg and 0.27mg/kg of Minnelide improved the percent of mice with tumors less than 1500mm^3^ tumor volume at 21 days (**Figure 4B**). Furthermore, treatment with 0.42mg/kg and 0.27mg/kg Minnelide resulted in a marked response in ∼61% (8/13) and ∼47% (7/15) of mice, respectively (**Figure 4C, D and E**), without any appreciable toxicity.

**Figure 4.**
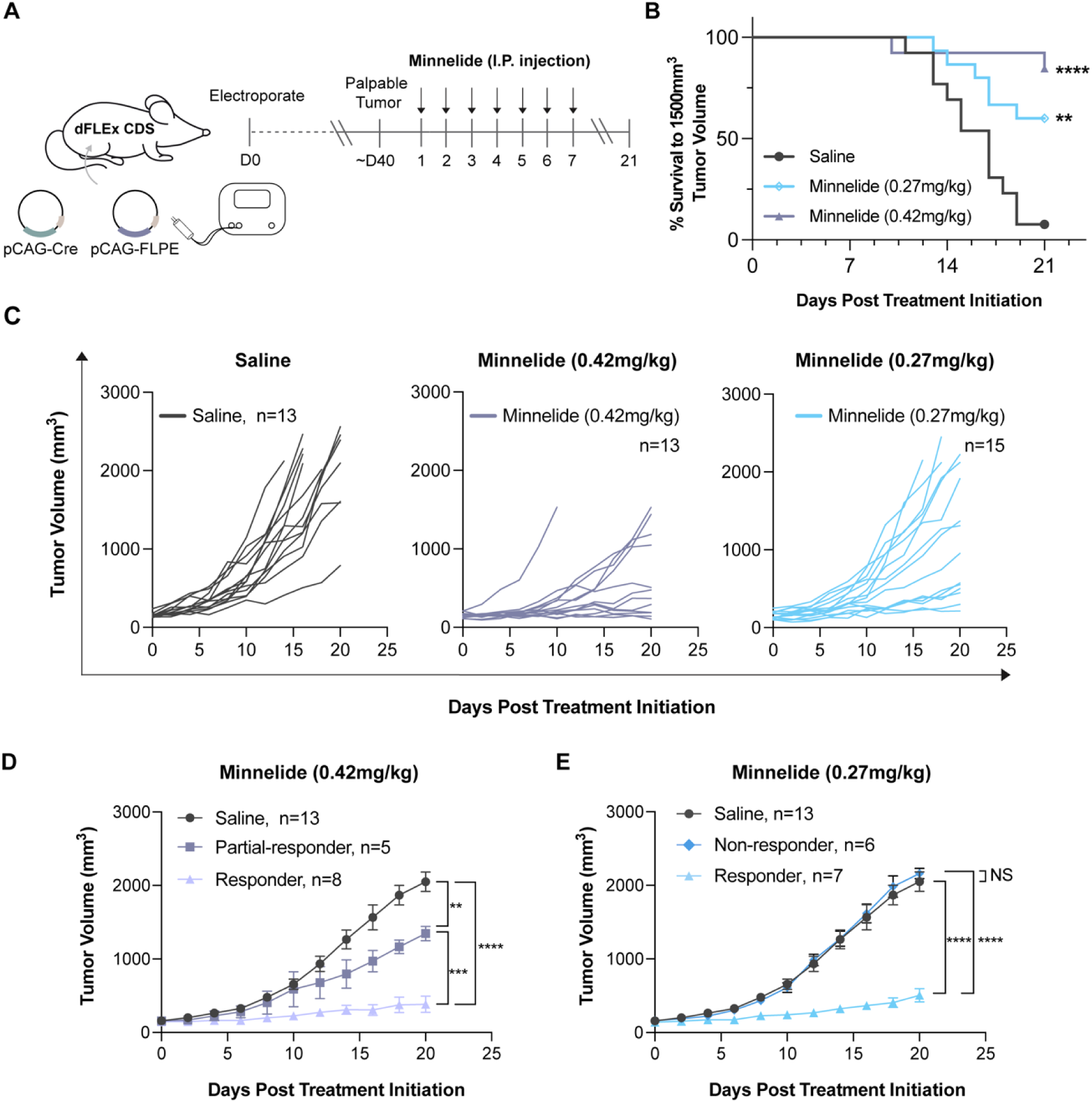
Minnelide reduces tumor growth in vivo in autochthonous sarcomas in dFLEX CDS mice. **A.** Schematic for dFLEx CDS tumor induction followed by Minnelide treatment in the dFLEx CDS model **B.** Kaplan-Meier curves indicating time to 1500mm^3^ tumor volume increased at both doses (0.42mg/kg and 0.27mg/kg) of Minnelide treatment **C.** Spider plots demonstrating tumor growth rates in control (saline), (0.42mg/kg), and (0.27mg/kg) Minnelide treated groups over 21 days **D.** Minnelide treatment with 0.42mg/kg resulted in reduced tumor growth in 8/13 mice while treatment with 0.27mg/kg resulted in reduced tumor growth in 7/13 mice. Data are presented as SEM; *p<0.05, **p<0.01, ***p=0.001, ****p<0.0001 by two-way ANOVA.

### Minnelide treatment significantly reduces tumor growth in human CDS xenograft models

To further substantiate the clinical potential of Minnelide in CDS, human CDS cells (Kitra-SRS and ECD1) were subcutaneously transplanted into NOD-scid IL2Rγnull (NSG) mice. Once tumors reached 200-300mm^3^, mice were treated with Minnelide via intraperitoneal injection once daily for 21 consecutive days. Dose selection was informed by the recommended starting dose and maximum tolerated dose reported in the Phase I Minnelide clinical trial for GI carcinoma (34). Treatment with both 0.21 mg/kg and 0.27 mg/kg of Minnelide, which approximates (44) to the human dose of 0.67 mg/m^2^ and 0.80 mg/m^2^ (**Figure 5A**), significantly reduced tumor growth of xenografts from both human CDS cell lines tested (**Figure 5B).** IHC on xenograft tumors demonstrated a reduction in Ki67 staining over the 21-day treatment period, and reduced p-RPB1 ser2/5 expression, indicating decreased RNAP II initiation and elongation, suggesting active transcription is impaired (**Figure 5C)**. Furthermore, Western blot analysis of xenograft tumors demonstrated a reduction in RPB1 expression (**Figure 5D)**. Consistent with previous studies (28,40,42) and our *in vitro* results (**Figure 3)**, these findings suggest Minnelide treatment induces RPB1 degradation and inhibition of transcription in human CDS xenograft tumors.

**Figure 5.**
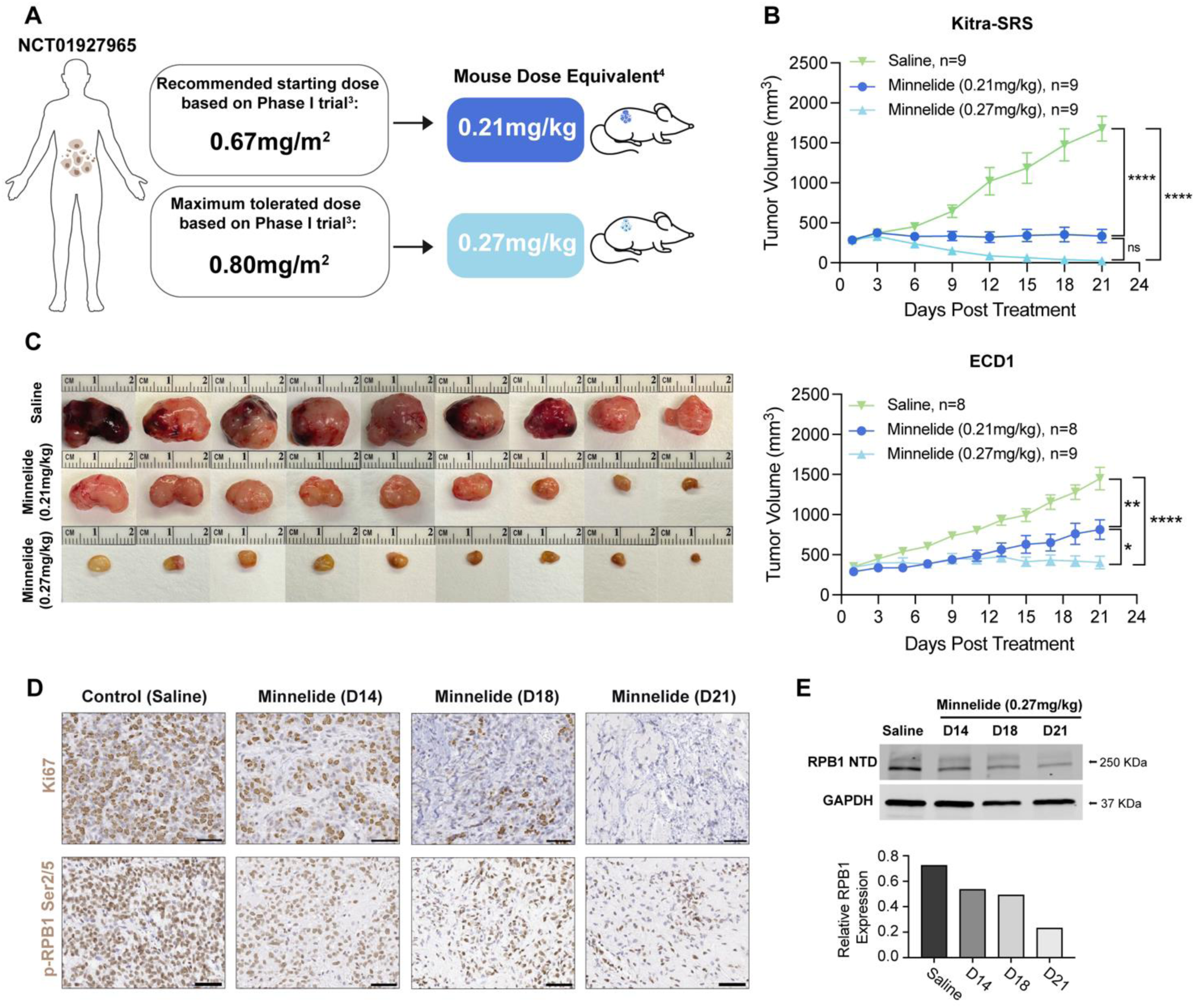
Minnelide reduces tumor growth in human CDS xenograft models. **A.** Dose selection was informed by the recommended starting dose and maximum tolerated dose reported in the Phase I Minnelide clinical trial for GI carcinoma (NCT01927965) 0.21 mg/kg and 0.27 mg/kg approximates the human dose of 0.67 mg/m^2^ and 0.80 mg/m^2^ respectively **B.** NSG mice were inoculated with Kitra-SRS or ECD1 human CDS cells then treated with Minnelide at 0.21 mg/kg or 0.27 mg/kg daily for 21 days. Tumor growth was significantly reduced in Minnelide-treated groups compared to controls (one-way ANOVA, *P < 0.05, ***P < 0.001, and ****P < 0.0001 at Day 21). **C.** Gross morphology of Kitra-SRS dissected tumors at humane endpoint or at 21 days of treatment. **D**. p-RPB1 Ser2/5 IHC on sections from saline and Minnelide treated Kitra-SRS CDS xenograft tumors at 14, 18 and 21 days of Minnelide treatment. **E.** Western blot of RPB1 expression in Kitra-SRS CDS xenograft tumors over a 21-day treatment period with 0.27mg/kg daily. Quantification of RPB1 expression relative to GAPDH loading control.

## Discussion

CDS is an aggressive sarcoma, which is driven by a neomorphic transcriptional activator, CIC::DUX4. Patients with CDS do not currently have effective systemic treatment options. A significant limitation to identifying and testing alternative treatment strategies has been the lack of robust immunocompetent pre-clinical models that recapitulate all stages of CDS tumor progression including tumor initiation within an intact immune system. Our previous attempts to generate a conditional CDS GEMM regulated by Cre recombinase were limited because of spontaneous sarcoma development in the absence of Cre (21). Here, we overcome this problem by employing a dual recombinase FLEx switch strategy which prevents spontaneous tumor formation. The observation that a dFLEx strategy is required to prevent spontaneous CDS tumor formation indicates that two frequently employed *loxP*-STOP-*loxP* cassettes, which we used in 3 previous CDS GEMM models that spontaneously developed tumors (21), is insufficient to completely prevent expression of an oncogene. Taken together, our results support the notion that *loxP*-STOP-*loxP* cassettes can, in rare instances, undergo spontaneous deletion or recombination in the absence of Cre recombinase. Whereas expression of an oncogene in a limited number of cells is typically insufficient to initiate tumorigenesis, the exceptional potency of CIC::DUX4 poses an extreme challenges for achieving controlled activation in vivo. This finding has important implications for modeling cancer in mice using *loxP*-STOP-*loxP* cassettes. Even when regulating less potent oncogenes or other neoantigens that cannot initiate cancer in the absence of Cre, scientists should be aware that low level gene expression of a conditional allele in the absence of Cre may occur. In this scenario, expression of the conditional allele may be sufficient to engage immune tolerance mechanisms that might subsequently impact the interaction of the immune system with tumor-initiating cells once Cre is expressed.

After electroporation of pCAG-Cre and pCAG-FLPE plasmids into the muscle of the hindlimb of dFLEx CDS mice, spatially and temporally controlled tumors arise that mimic human CDS by histology, immunohistochemistry, and expression of CIC::DUX4 transcriptional targets. Using a small molecule library screen, we also identified Minnelide, an inhibitor of RNA polymerase II (RNAP II)-mediated transcription, as a potential therapeutic approach to CDS. Minnelide induces apoptosis in human CDS cells and inhibits the growth of both dFLEx CDS autochthonous tumors and human CDS xenograft tumors *in vivo.* Considering Minnelide has already been tested in Phase I and II clinical trials for advanced gastrointestinal carcinoma and pancreatic cancer (34,35), and is currently being studied in ongoing clinical trials of gastric cancer (NCT05566834) and small cell lung cancer (NCT05166616), our results highlight the potential for Minnelide as a the treatment for CDS.

Previous studies have shown that Minnelide is rapidly converted to Triptolide when exposed to phosphatases in the blood and/or tissues (33). Triptolide then directly binds XPB, inhibiting its ATPase function to cause stalling of RNAP II at gene promoters (28,40,42,45). Prolonged stalling of RNAP II leads to hyperphosphorylation of RPB1 (the largest subunit of RNAP II) at Ser5, then subsequently, ubiquitination and proteasomal mediated degradation (28,40,42,45). Consistent with this mechanistic model, we show that human CDS cells treated with 10 μM Minnelide demonstrate a transient increase in RPB1 phosphorylation at Ser5 and near-complete degradation of RPB1 by 120 minutes. Furthermore, co-treatment with proteosome inhibitor Epoxomicin or expression of an XPB mutant (42) (XPB C342T) rescued RPB1 degradation. These results suggest that, similar to other cancer types, Minnelide directly targets XPB in CDS cells, resulting in proteasomal-mediated degradation of RPB1 and impaired RNAP II-mediated transcription.

Following RPB1 degradation, there are several potential pathways to induce cell death. One possibility is that inhibition of transcription leads to widespread mRNA decay over time with progressive loss of essential proteins, which results in a passive form of cell death termed accidental cell death (46). A second possibility is that inhibition of transcription leads to the loss of expression of essential genes that are specifically regulated by CIC::DUX4, similar to G3 MB, where Minnelide treatment decreases *MYC* expression (28). Alternatively, a recent elegant study demonstrated that degradation of RPB1 itself can trigger apoptosis through a transcription-independent mechanism, termed the Poll II degradation-dependent apoptotic response (47). In this that study, RBP1 mutants were used to uncouple RNAP II degradation from its transcriptional activity, revealing that RNAP II degradation alone is sufficient to induce apoptosis independent of mRNA decay. Instead, degradation of RNAP II shifted the localization of BCL2L12 from the nucleus to mitochondria via a polypyrimidine tract-binding protein 1 (PTBP1) mechanism to trigger the intrinsic pathway of apoptosis (47). This mechanism of cell death in CDS would provide a rationale for the specificity of Minnelide in CDS cells over many normal adult tissues, whose mitochondria are often refractory to pro-apoptotic signaling (48).

Considering CDS is driven by the activation of a unique oncogenic transcriptional program, we anticipated Minnelide would suppress this CIC::DUX4-specific transcriptional program, resulting in loss of essential gene expression after mRNA decay and induction of cell death. Unexpectedly, there were few CIC::DUX4 target genes whose expression decreased 2-24h following Minnelide treatment. Furthermore, while significant degradation of RPB1 and cell death are observed after 48h of Minnelide treatment, a substantial number of transcriptional changes still occur even after 72h of Minnelide treatment. These results together with recent findings that RNAP II degradation in the absence of mRNA decay is sufficient to trigger the intrinsic pathway of apoptosis (47) suggest that apoptosis in CDS cells following Minnelide treatment may be triggered via the RNAP II degradation-dependent apoptotic response (47) in a manner independent of mRNA decay and changes in transcription.

As there is an urgent unmet clinical need for novel therapeutic strategies for CDS, we used two different murine models to test the efficacy of Minnelide against CDS *in vivo*. We started by inducing tumors in the dFLEx CDS model, then treated mice daily for 21 days with either 0.42mg/kg Minnelide, 0.27mg/kg Minnelide or saline. Interestingly, at 0.27mg/kg Minnelide there were responders and non-responders in the dFLEx CDS model. In contrast, the human CDS xenograft model demonstrated reduced tumor growth in almost every mouse tested.

These results not only highlight the value of the dFLEx CDS model, but also emphasize the importance of utilizing multiple pre-clinical models when testing potential therapies for CDS. Future studies will be performed to explore the mechanisms driving response vs non-response in the dFLEx model, which might be relevant in future clinical trials of Minnelide in CDS patients.

## Methods

### Sex as a Biological Variable

All *in vivo* studies included both male and female animals, and similar results were observed in both sexes.

### Study approval

All animal experiments were approved by both Duke University Animal Care and Use Committee (protocol number A014-22-01) and by the University Health Network’s Princess Margaret Cancer Centre Animal Care Committee, aligned with guidelines from the Canadian Council on Animal Care (protocol number 6825).

### Generation of Transgenic Animals

dFLEx CDS mice were generated by Ozgene (Perth, WA, Australia). An inverted 3xHA CIC exon and inverted CIC/Dux4-CTD exon (inverted 3xHA_CIC exon_lox2272_loxP_FRT’_FRT_inverted CTC_Dux4-CTD exon_FRT’_SV40pA_KT3_FRT_DUX4-CTS exon_SP6 promoter_BGHpA) was cloned into a Rosa26 targeting construct. After sequence verification, the construct was electroporated into ES cells and selected in antibiotic (neomycin) containing media. Clones containing the knock-in were screened and validated by qPCR, then injected into goGermline blastocysts which were transplanted into female nurse mice for gestation and delivery of goGermline heterozygous males. GoGermline heterozygous male mice were then bred to generate 100% ES cell derived dFLEx mice.

### Genotyping

Genomic DNA (gDNA) was purified from tail clips using Qiagen DNeasy Blood and Tissue kit (Cat. 69504). Primers designed to amplify across a region of the stop cassette into the inverted exon 1 (**Supplementary Table 1**) were used to validate the presence of the dFLEx allele. PCR was performed using 2X Taq FroggaMix (Cat. FBTAQM) and optimized for amplicon size. To validate recombination, gDNA was purified from dFLEx tumors, and primers were designed to amplify across the LoxP and FRT sites (**Supplementary Table 1**). PCR on dFLEx tumors was performed using 2X Taq FroggaMix. As a control, PCR utilizing the same primers was conducted on tail clips from dFLEx mice. To accommodate the unrecombined large amplicon size, PCR was performed using NEB LongAmp Taq DNA Polymerase (Cat. M0323S).

### dFLEx CDS tumor formation and evaluation of Minnelide in vivo

6-12 week old dFLEx CDS mice were anaesthetized, then 30ug of naked pCAG-Cre (Addgene #13775) and pCAG-FLPE (#13787) plasmid DNA diluted in sterile saline was injected into the gastrocnemius using a 31-gauge insulin syringe. As previously described (49), needle electrodes with a 5 mm gap were inserted into the muscle to encompass the DNA injection site, and electric pulses were delivered using an electric pulse generator (Electro Square Porator ECM830; BTX, San Diego, CA). Three 100 V pulses followed by three additional 100 V pulses of the opposite polarity were administered to each injection site at a rate of 1 pulse per 50 ms with each pulse being 200 ms in duration. Palpable tumors were detected starting at 30 days post electroporation. For the evaluation of Minnelide in the dFLEx CDS model, mice were treated daily for 21 consecutive days with 0.42 mg/kg Minnelide, 0.27 mg/kg Minnelide, or saline by intraperitoneal injection, beginning at the time of tumor palpation. Tumors were measured every 2 days by digital caliper, and tumor volume was calculated using the formula (length x width^2^)/2. Tumors 170256 and 170220 exhibited intermediate tumor size between responder and non-responder groups, and were therefore excluded from the analysis in Figure 4E. pCAG-Cre and pCAG-FLPE were a gifts from Connie Cepko (Addgene plasmid # 13775 and # 13787)(50). Control sarcomas from KRAS^G12D^; p53^fl/fl^ (KP) mice were generated as previously described (51).

### Cell Culture

DMEM (Cat. 11995065), Fetal bovine serum (FBS, Cat. A3160702) and Penicillin/ Streptomycin (P/S, Cat. 15140122) were purchased from GIBCO, RPMI was from Sigma-Aldrich (Cat. R8758). Kitra-SRS (CIC::DUX4 sarcoma cell line (17)), TOPCDS (mouse CIC::DUX4 sarcoma cell line (21)), A-673 (Ewing sarcoma cell line, ATCC), and WT MEFs were cultured in DMEM/10% FBS/P/S. ECD1 (CIC::DUX4 sarcoma cell line (7)), X1C1 (CIC::DUX4 sarcoma cell line(18)), CDS2 (CIC::DUX4 sarcoma cell line (52)), and Rh-4 (Fusion-positive Rhabdomyosarcoma cell line) were cultured in RPMI/20% FBS/P/S. All cells were cultured at 37 °C in a 5% CO_2_ atmosphere.

### Xenograft tumor formation and evaluation of Minnelide in vivo

6-8 week old NOD-scid IL2Rγnull mice (The Jackson laboratory, 005557) were subcutaneously transplanted with 1.0x10^7^ Kitra-SRS or ECD1 cells resuspended in a 100μl, 1:1 mixture of culture media and Matrigel (Corning, 356231). Palpable tumors at the site of injection were detected starting at day 7 after transplantation. After tumors reached ∼200-300mm^3^ by digital caliper measurement using the formula (length x width^2^)/2, mice were randomized into groups (0.21mg/kg Minnelide, 0.27mg/kg Minnelide, and saline). Minnelide was initially dissolved in H_2_O, then further diluted in saline. Each mouse received 0.21mg/kg Minnelide, 0.27mg/kg Minnelide or saline by intraperitoneal injection daily for 21 consecutive days. Tumor volume was measured every 2-3 days by digital caliper, and tumor volume was calculated using the formula (length x width^2^)/2. Because individual mice reached ∼200-300mm^3^ tumor volume at different times, tumor measurements obtained within each 3-day interval were averaged and assigned to the midpoint of that interval (e.g., day 2), allowing consistent comparison of growth kinetics across animals.

### Western blot

Cells lines were maintained in standard growth media until 80% confluency. Using Trypsin-EDTA, the cells were lifted, collected in Phosphate-Buffered Saline (PBS), then pelleted by centrifugation (300xg for 3 minutes). Lysates were made using Pierce RIPA buffer (Thermo Fisher Scientific, Cat. 89900) supplemented with 1% SDS, Halt protease inhibitor (Thermo Fisher Scientific, Cat. 78441), Benzonase, and PhosSTOP phosphatase inhibitor (Roche, Cat. 4906845001), then protein was quantified using Pierce BCA protein assay kit (Thermo Fisher Scientific, Cat. 23225). Heat-denatured proteins were loaded onto a 4-20% Tris-glycine gel, run at 100v in 1x Tris/Glycine/SDS buffer, and then wet-transferred at 350mA for 1 hour at 4 °C. When detecting the CIC::DUX4 fusion, all above steps were completed in a single day due to the unstable nature of the fusion protein. CIC::DUX4 (∼260kD) was probed using anti-DUX4 antibody (Abcam, ab124699, 1:1000) and anti-HA antibody (Cell signaling, 3724, 1:1000). RPB1 and p-RPB1 ser2 and ser5 were detected using anti-RPB1 (Cell signaling, 54020, 1:1000) and anti-p-RPB1 antibody (Cell signaling, 54020, 1:1000) with GAPDH (Cell signaling, 2118, 1:4000) or β-actin (Cell signaling, 3700, 1:4000) as a loading controls. Images were acquired on a LI-COR Odyssey CLx and processed using the Image Studio Software.

### Immunohistochemistry

As previously described (21) tissue samples were fixed in 10% formalin/70% EtoH, then embedded in paraffin blocks. 0.4μM sections were mounted and stained with hematoxylin and eosin (H&E) or antibodies. Detection of antibodies was performed using the Vectastain Elite ABC-HRP Kit (Vector Labs, VECTPK6100) and DAB Peroxidase Substrate Kit (Vector Labs, VECTSK4100). Antigen unmasking was performed with a citrate buffer pH 6.0 by a modified microwave retrieval method followed by boiling. Images were captured on the Aperio AT2 brightfield scanner (Leica Biosystems) using the UPlanS Apo 20x / 0.75 NA (high-resolution 20x objective). The following antibodies were used for immunohistochemistry: HA (Cell Signaling, 3724) at 1:800, ETV4 (Proteintech 10684-1-AP) at 1:1800, CD99 (Thermo-Fisher, MA5-12287) at 1:200, WT1 (Thermo-Fisher, MA5-32215) at 1:2000, Pan-cytokeratin (abcam, ab9377) at 1:200, Desmin (abcam, ab15200) at 1:200 and Phospho-Rpb1 CTD (Ser2/Ser5) (Cell Signaling, 13546) at 1:100.

### RNA Sequencing

RNA was extracted and purified from flash frozen tumors and cells using a Qiagen RNeasy Plus kit (Cat. 74134). As described previously(21), high quality RNA (RIN >7) was divided into duplicate from which 150bp paired-end, rRNA-depleted, libraries were made using the Illumina TruSeq RNA library Prep Kit (Illumina, CA, USA). Libraries were quantified using the KAPA Library Quantification kit (KAPA Biosystems, MA, USA), multiplexed, clustered onto flowcells, and then sequenced using an Illumina HiSeq 4000 sequencer (or equivalent platform) by GENEWIZ (Azenta, NJ, USA). Raw sequencing reads were trimmed using Trimmomatic v0.39 (ILLUMINACLIP:TruSeq3-PE-2.fa:2:30:10:2:keepBothReads LEADING:3 TRAILING:3 MINLEN:36) and then aligned to the mm39 reference genome (dFLEx) or hs38 using default parameters in STAR v2.7.10a. FeatureCounts (Subread v2.0.3) was used to compile a count table from sorted and indexed BAM files which was loaded into DESeq2 to calculate differential expression. Gene set variation analysis (GSVA) was used to visualize the top enriched pathways in dFLEx CDS tumors, while the *fgsea* R package was used to identify significantly enriched pathways in the Minnelide time-course RNA-seq dataset from Kitra-SRS cells. To assess how Minnelide might induce apoptosis, genes were curated from the GSEA HALLMARK_APOPTOSIS gene set, and manually assigned to either intrinsic apoptosis pathway or the extrinsic apoptosis pathway. Differential expression results were used to evaluate the expression of the apoptosis associated genes at 12h, 24h, 48h and 72h compared to the non-treated control cells. Statistical significance was defined as padj < 0.05. Genes with padj < 0.05 and log2FC > 0 were classified as “Upregulated,” padj < 0.05 and log2FC < 0 as “Downregulated,” and all others as “Not Significant.”

### Cell Viability Drug Screen

The Tocriscreen Epigenetics 3.0 compound library (Cat. 7578) was used in this screen. Kitra-SRS, CDS#2 and X1C1 CDS cell lines were plated at a density of 3-5 x 10^4^ cells per well in 96-well plates. In technical triplicate, cells were incubated in drug at four different concentrations (1, 10, 100, and 1000nM) for 72 hours and then assayed using Cell Titer Glo luminescence (Promega, G7572) to assess cell viability. The plates were read using a CLARIOstar Plus plate reader, and RLU was normalized to DMSO-treated cells. For visualization, 20 compounds with the largest overall effect size and variance were selected. Compounds were ordered by drug class, and percent viability across conditions was plotted as a heatmap using the pheatmap package in R.

### Cell Viability (Cell Titer Glo) assay

Cell lines were were plated at a density of 3-5 x 10^4^ cells per well in 96-well plates. In technical quadruplicates, cells were incubated in Minnelide (dissolved in H_2_O) at concentrations ranging from 0nM to 200nM for 48h. Cell viability was assayed using Cell Titer Glo luminescence (Promega, G7572) to assess cell viability. The plates were read using a CLARIOstar Plus plate reader, and RLU was normalized to 0nM untreated cells.

### Flow Cytometry

Dead Cell Apoptosis Kit with Annexin V for Flow Cytometry (Thermo-Fisher, V13245) was used for the detection of apoptotic cells. Kitra-SRS and ECD1 cells were Trypsinized then pelleted by centrifugation (300xg for 3 minutes). Cell pellets were resuspended in 1X Annexin-binding buffer, passed through a 70 μm cell strainer, and then transferred to Falcon round-bottom polystyrene test tubes equipped with 35 μm nylon mesh caps. Cell suspensions were adjusted to a density of 1.0 × 10⁶ cells/mL, then Alexa Fluor 488 Annexin V and Propidium iodide were added according to the manufacturer’s instructions. Cells were incubated for 15 minutes at room temperature, then analyzed on a BD LSR Fortessa X20 flow cytometer.

### Immunofluorescence

Cells were grown on 12 mm poly-L-lysine coated glass coverslips (Neuvitro, Cat # GG-12-15-PLL), washed with Dulbecco′s Phosphate Buffered Saline with MgCl₂ and CaCl₂ (Sigma, D8662) for 5 min at 37°C, prior to fixation in 4% PFA (Electron Microscopy Sciences, Cat #15710) for 10 min at room temperature (R.T.). Coverslips were incubated in blocking buffer (0.2% Triton-X-100, 2.5% BSA) for 1h at R.T, then in Cleaved-caspase 3 primary antibody solution (Cell signaling, 9661, 1:400) overnight at 4°C. Next, coverslips were incubated with goat anti-rat secondary antibody for 1h at R.T. (Alexa Fluor 488, 1:1000,Invitrogen Cat# A-11001). Coverslips were mounted on SuperFrost slides (Fischer Scientific, Cat #22-037-246) with ProLong Gold anti-fade Mountant with DAPI DNA Stain (Invitrogen, Cat #P36931). Images were obtained on the Leica Stellaris confocal microscope, equipped with an HC PL APO HCX PL APO 40x/0.95 NA CORR (0.11 - 0.23mm coverslip correction) lens, using a Hamamatsu Camera, and processed using Fiji.

### EdU Incorperation assay

ECD-1 cells were grown to 70% confluence on 12 mm poly-L-lysine coated glass coverslips (Neuvitro, Cat # GG-12-15-PLL) and treated with Minnelide at 25nM concentration or vehicle for 48 h. 5-ethynyl-2’ -deoxyuridine (EdU) incorporation assay was carried out according to manufacturer’s instructions (Click-iT™ Plus EdU Cell Proliferation Kit for Imaging, Alexa Fluor™ 647 dye, Invitrogen, Cat # C10640). Briefly, cells were incubated with 10 µM EdU for 1 h prior to fixation in 3.7% PFA for 15 min at R.T. Following this, the cells were washed in 3% BSA and permeabilized using 0.5% Triton X-100 for 20 min at R.T. Cells were then incubated in Click-iT detection reagents (Alexa Fluor 647, 1:1000) for 30 min in the dark at R.T, followed by nuclear staining using 1 ug/mL DAPI. Coverslips were mounted on SuperFrost slides (Fischer Scientific, Cat #22037246) using ProLong gold antifade mounting media (Invitrogen, Cat # P36930). All images were taken using a Leica Stellaris 5 confocal microscope equipped with an HC PL APO 20x/0.75 NA CS2 objective. Frequency of EdU positive cells was calculated on Fiji using manual thresholding parameters.

### Generation of XPB mutant

The XPB C342T mutant construct was generated by Genescript. For Lentiviral production, 293FT cells were transfected with XPB_Cys342Thr_pGenLenti, pRSV-Rev, PMDLg/pRRE and pMD2.G using Lipofectamine 2000 (Thermo Fisher Scientific). Kitra-SRS and ECD1 cells were transduced with viral supernatant using a concentration of 2 ug/mL polybrene, followed by antibiotic selection using 0.25-0.5mg/mL Puromycin.

### Statistics

Results are presented as means ± s.e.m. unless otherwise indicated. Prior to statistical analysis, all datasets were displayed graphically, (box-and-whisker in GraphPad Prism 10), and assessed for normality using the Shapiro–Wilk test to determine whether parametric or non-parametric tests should be used. One-way ANOVA (at day 21) or Two-way ANOVA was used to compare tumor growth between Minnelide and Saline treated groups. Survival data were analyzed using Kaplan-Meier curves with the log-rank test for statistical significance. All calculations were performed using Prism 10 (GraphPad). Additional details on each statistical test used can be found in the Supporting Data Values file.

## Data Availability

All RNA-sequencing data will be freely accessible through the Gene Expression Omnibus (GEO). The dFLEx CDS mouse line will be available at The Jackson Laboratory (Stock no. 041280). Values for all data points in graphs are reported in the Supporting Data Values file.

## Author Contributions

This project conceptualized and designed by P.G.H, D.G.K and M.R.B with support from K.M.O and J.E.H. All experiments were performed by M.R.B with support from P.G.H, A.V.S, and R.B. Animal breeding and maintenance was performed by B.A. The paper was written by M.R.B with editing from P.G.H and D.G.K. All authors approved the submission of this manuscript.

## Acknowledgements

We would like to thank Roger Askew and the members of Ozgene for their help with generation of the dFLEx CDS mouse model and Benjamin Alman for his mentorship and support of M.R.B. We thank Satoshi Takenaka from Osaka University Graduate School of Medicine for providing the human CDS cell line Kitra-SRS, Takuro Nakamura from Japanese Foundation for Cancer Research for providing the human CDS cell line ECD1, and Tadashi Kondo from National Cancer Center Research Institute for providing the human CDS cell lines NCC-CDS-X1C1 and NCC-CDS2-C1.

