## Supplemental figures for "Efficacy of Minnelide in a Next-Generation Dual-Recombinase Regulated Genetically Engineered Mouse Model of CIC::DUX4 Sarcoma"

#### Supplementary Figure 1

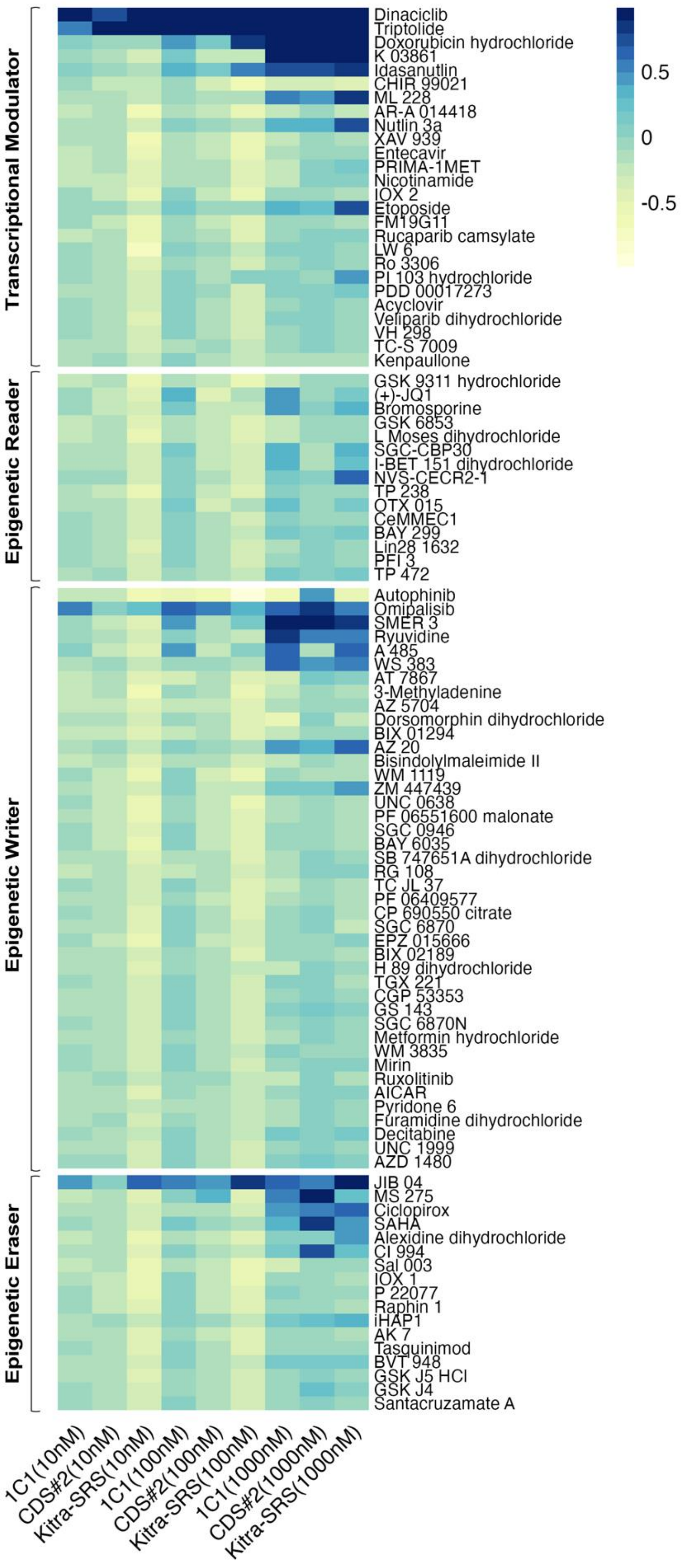


**Figure S1.** Viability drug screen on human CDS cells using the Tocriscreen Epigenetics 3.0 compound library. Top 100 compounds at 10nM, 100nM and 1000nM concentrations.

### Supplementary Figure 2

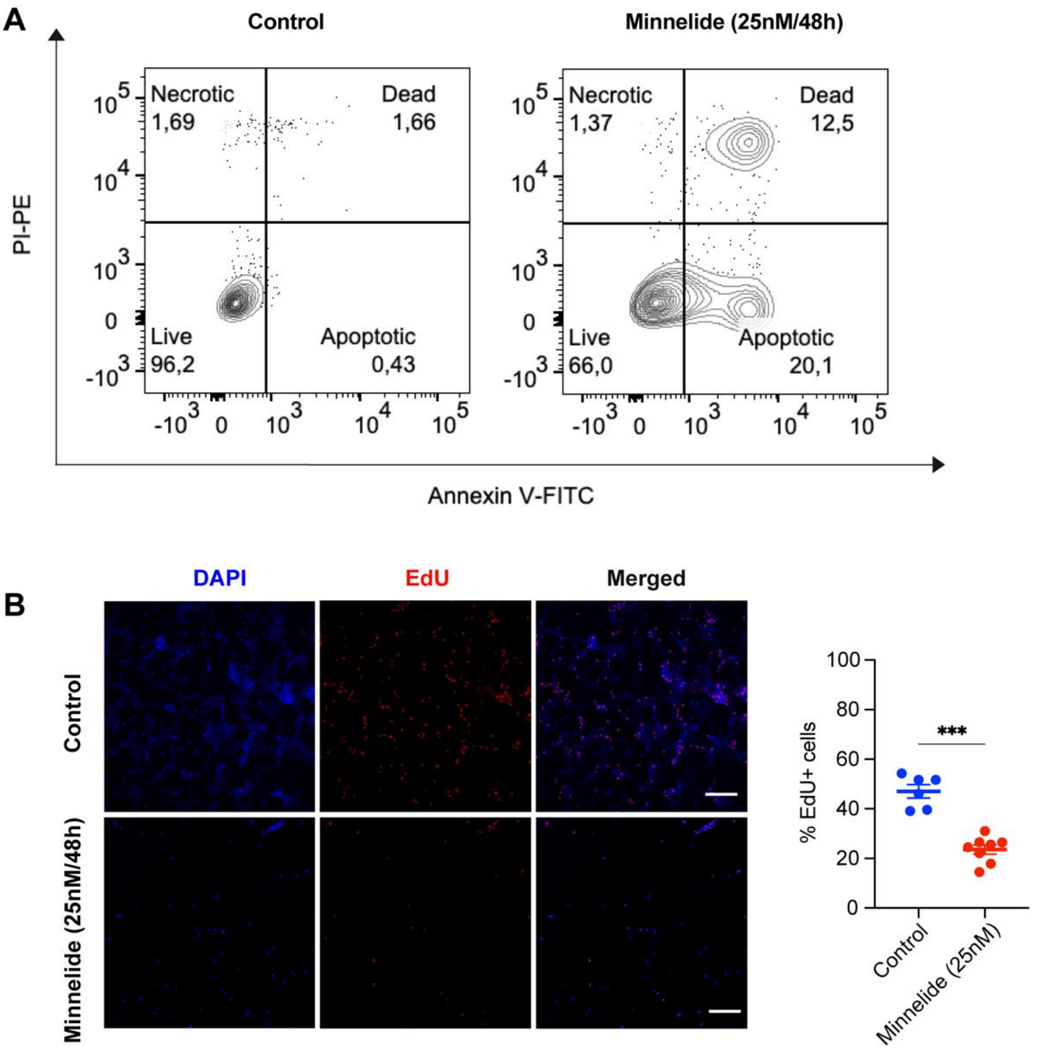


**Figure S2. A.** Annexin V/PI flow cytometry on ECD1 CDS cells treated with Minnelide for 48h. **B.** EdU labeling on human ECD1 CDS cells treated with Minnelide for 48h. ***P ≤ 0.001.

### Supplementary Figure 3

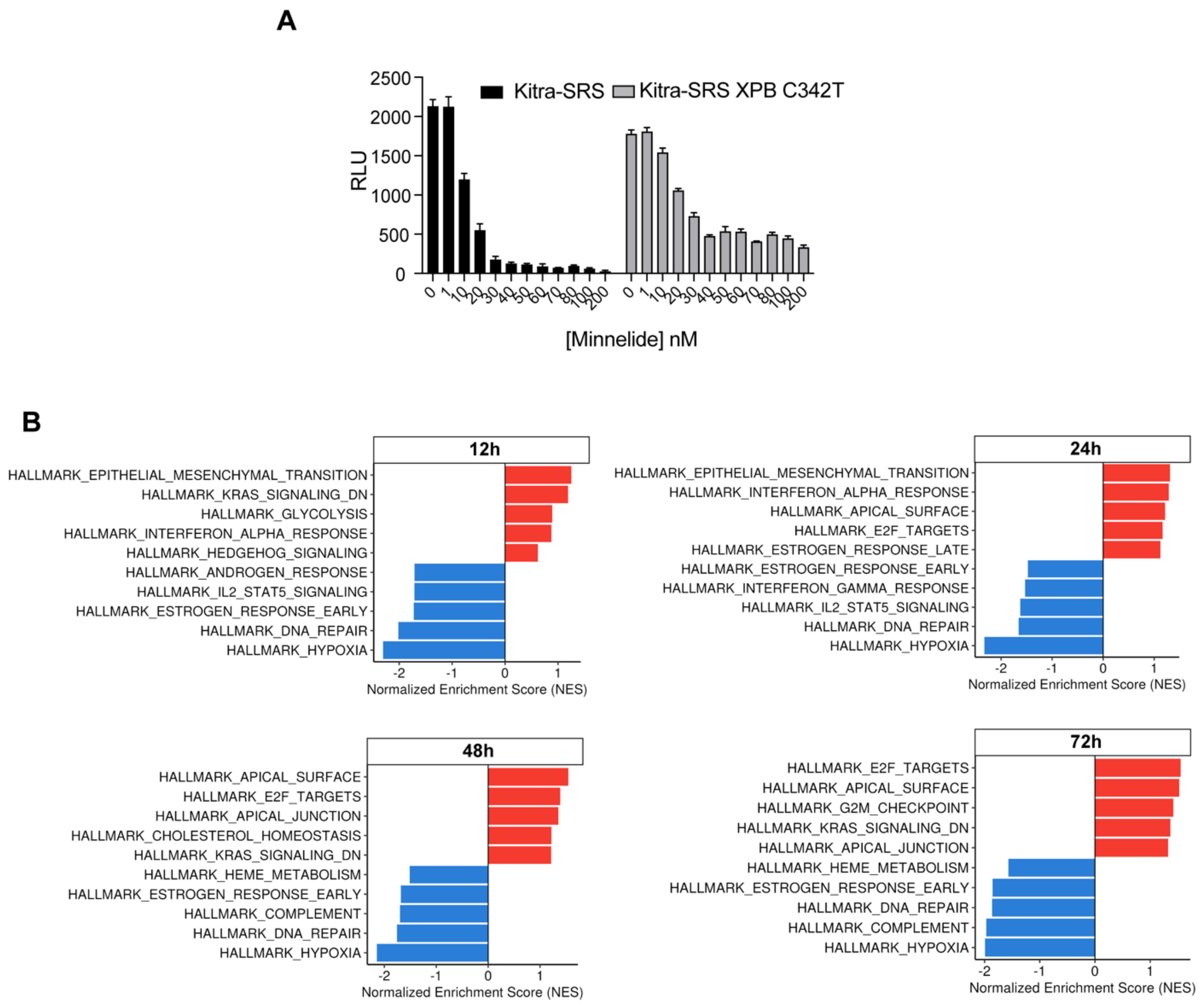


**Figure S3. A**. Cell-titer-glo assay demonstrates human CDS (Kitra-SRS) cells expressing XPB C342T are resistant to Minnelide. **B.** GSEA pathway analysis on Kitra-SRS cells treated with 25nM Minnelide for 12h, 24h, 48h and 72h

#### Supplementary Figure 4

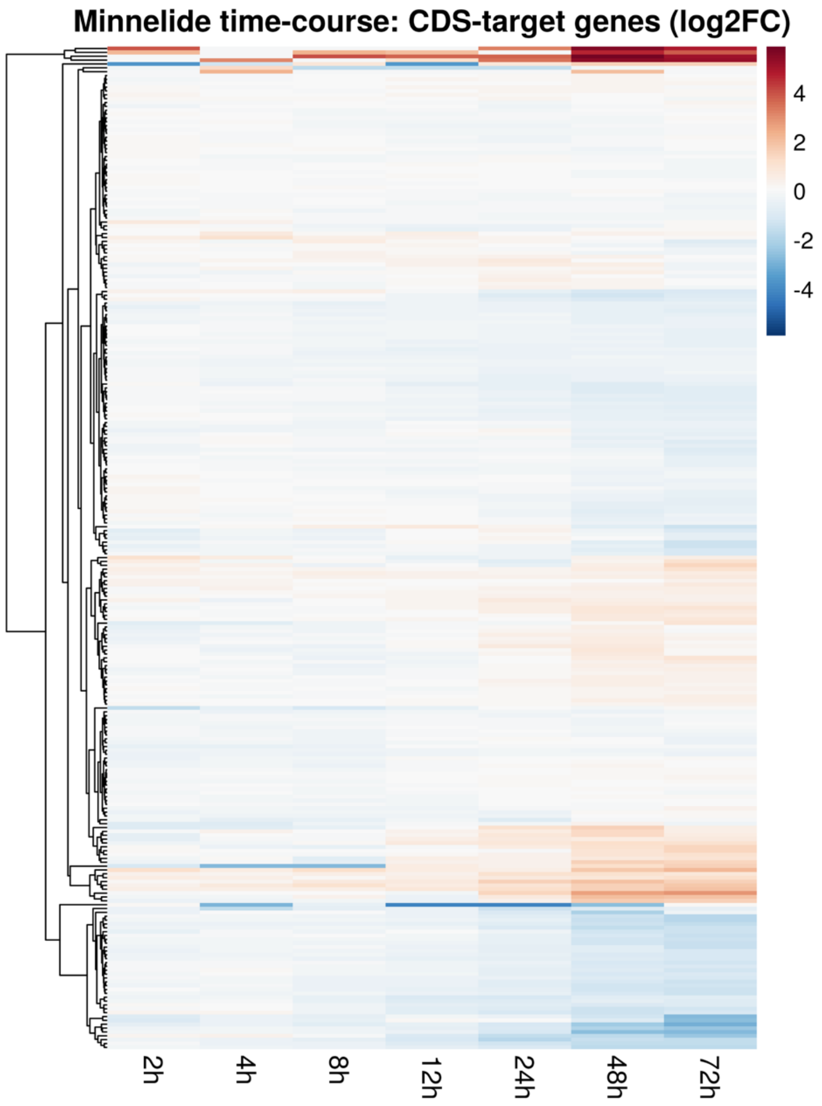


**Figure S4.** Heatmap showing expression of CIC::DUX4 target genes following Minnelide treatment for 2, 4, 8, 12, 24, 48, and 72 hours. The majority of CDS target genes, including *ETV1/4/5*, remained largely unchanged until 72 hours post-treatment.
